# A cross-sectional survey of Social Media Anxiety among students of University of Nigeria

**DOI:** 10.1101/666701

**Authors:** Deborah O. Aluh, Thelma Chukwuobasi, Adaobi U. Mosanya

## Abstract

**Background:** Social anxiety is one of the most prevalent and disabling anxiety disorders with lifetime prevalence rates ranging from 2 to 16% s in different populations. Considering the rising use of social media among university students, it is necessary to assess their social anxiety as a result of the use of social media platforms since social anxiety can affect social interaction in social media

**Methods:** The current study employed a cross-sectional descriptive approach and was carried out among undergraduate students of University of Nigeria, Nsukka. The Social Anxiety Scale for Social Media Users (SAS-SMU) which is a data collection tool to assess levels of social anxiety experienced by university students while using social media platforms was used in the current study. Data were analyzed with IBM Statistical Products and Service Solutions (SPSS) for Windows, Version 20.0.

**Results:** A total of 228 out of the 380 questionnaires distributed were filled and returned (60% response rate). Social media usage was highest for WhatsApp (4.4±0.902) and Facebook (3.3±1.055). Social media anxiety was higher in females (69.00±12.59) than males (68.42±12.06) although this difference did not reach statistical significance (t = −0.356, p = 0.864). Social media usage was higher in females (35.02±5.04) than males (34.58±6.01) but the difference did not reach statistical significance (t = −0.603, p = 0.314). There was a non-significant negative association between Social media usage and social media anxiety (r = –0.051, p = 0.4450). More than half of the students (55.7%, n = 127) had social media anxiety.

**Conclusion:** In conclusion, there was a negative non-significant correlation between social media usage and social anxiety. Investigations regarding social anxiety in social media are scarce from low and middle income countries and this is the first from an African country.

## Introduction

Anxiety is currently the second most burdensome psychiatric disorder with the burden peaking at early adulthood^1^. Social anxiety is one of the most prevalent and disabling anxiety disorders with lifetime prevalence rates ranging from 2 to 16% s in different populations^2,3^. In Nigeria, the lifetime prevalence social anxiety among university students was found to be 9.4% which is comparable to findings from other parts of the world^4^. Disability caused by anxiety disorders has been reported to be akin to chronic physical ailments such as arthritis, hypertension and diabetes; anxiety disorders usually occur earlier in life and consequently, have a longer duration of disability ^5^.

Social anxiety has been defined as “a type of anxiety-related problem resulting from when people are fearful or anxious when interacting with or being negatively evaluated and scrutinized by other people during social interactions in a social setting” ^6^. Some factors have been implicated in the development of social anxiety. Weisman *et al.*, in their research found that social anxiety was correlated with low perceived intimacy and closeness in peer, friendship, and romantic relations^7^. Their study also discovered that people with social anxiety disorder thought of themselves as inferior, having low social rank, and behaved submissively. Many studies have shown an association between people’s behaviors in social media space and anxiety. Social connectedness as a result of Facebook use has been shown to be negatively correlated with anxiety in a study by Grieve and his colleagues ^8^. In the process of assessing the association between internet use and internalizing problems, the use of the internet for purposes apart from communication was found to be correlated with depression and social anxiety^9^. Sedentary computer has also been shown to lead to depressive and anxiety disorders ^10^.

Concerns about privacy emerging in social media platforms have been a subject of interest in recent times, since they’ve been shown to affect social anxiety^11^. Privacy concerns comprise potential privacy risks about personal information or distinguishing characteristics (e.g., unintentional disclosure of private comments or messages, personal information including birthday, home address, mobile phone numbers and personal photographs) revealed through social media platforms^12^. A study by Liu and his colleagues in 2013 showed that concerns about privacy had an impact on social anxiety ^13^. Considering the rising use of social media among university students, it is necessary to assess their social anxiety as a result of the use of social media platforms since social anxiety can affect social interaction in social media ^11^.

There’s sparse literature on the association between social media and anxiety. Most of the available data originate from high-income countries^2^. This study aimed to investigate University students’ social anxiety when using social media in Nigeria.

## Methods

The current study employed a cross-sectional descriptive approach and was carried out among undergraduate students of University of Nigeria, Nsukka. University of Nigeria is one of the premiere government owned universities in Nigeria created in 1970 and boasts of having students with diverse ethnicities and backgrounds. Given that a total 28047 students were enrolled in the university as at September 2018, allowing 0.5% margin of error at 95% confidence interval, the minimum sample size was calculated to be 379. Students who were willing taking part in the survey were conveniently sampled in their different faculties. The questionnaires were distributed to all consenting students who were present in their faculty lecture theatres during the study period.

A cover letter was attached to the survey instrument which informed respondents of the purpose of the survey and assured them of confidentiality and anonymity. Ethical clearance for the study was gotten from the University of Nigeria Ethical committee.

## Data Collection and Instrument

The Social Anxiety Scale for Social Media Users (SAS-SMU)^11^ which is a data collection tool to assess levels of social anxiety experienced by university students while using social media platforms was used in the present study. The 35-item questionnaire had three main sections. The first part solicited information on respondents’ gender, age, department, and education level. In the second part, respondents were asked questions on how often they used selected social media platforms on a 5-point Likert-type scale ranging from 1-never to 5-always. The third part assessed Social anxiety by items on shared content anxiety, privacy concern anxiety, interaction anxiety, and self-evaluation anxiety on a 5-point Likert scale ranging from 1-strongly disagree to 5-strongly agree. The questionnaire was pre-tested using 15 pharmacy students and had good internal consistency with a Cronbach alpha of 0.87. The data obtained during the pre-test were not included in the study data.

## Data analysis

Data were analyzed with IBM Statistical Products and Service Solutions (SPSS) for Windows, Version 20.0. Descriptive statistics such as means and standard deviation were used to characterize the data obtained. Pearson correlation was used to determine the association between social media usage and social media anxiety with significance set at <0.05. Independent sample t-test was used to compare means of continuous variables between genders. Socio-demographic variables were presented as frequencies, percentages.

## Results

A total of 228 out of the 380 questionnaires distributed were filled and returned (60% response rate). There were more female (57%, n = 130) than male respondents in the present study. Almost three-quarters of the respondents were mostly between the ages of 18 and 24 years (71.5%, n = 163). The respondents were mostly in their fourth year of study (29.8%, n = 68) and from the faculty of pharmaceutical sciences (54.4%, n = 124). (Table 1) Social media usage was highest for WhatsApp (4.4±0.902). Skype and LinkedIn were the least frequently used social media platforms by the respondents (1.4±0.646, 1.4±0.805) respectively. (Table 2)

**Table 1:**
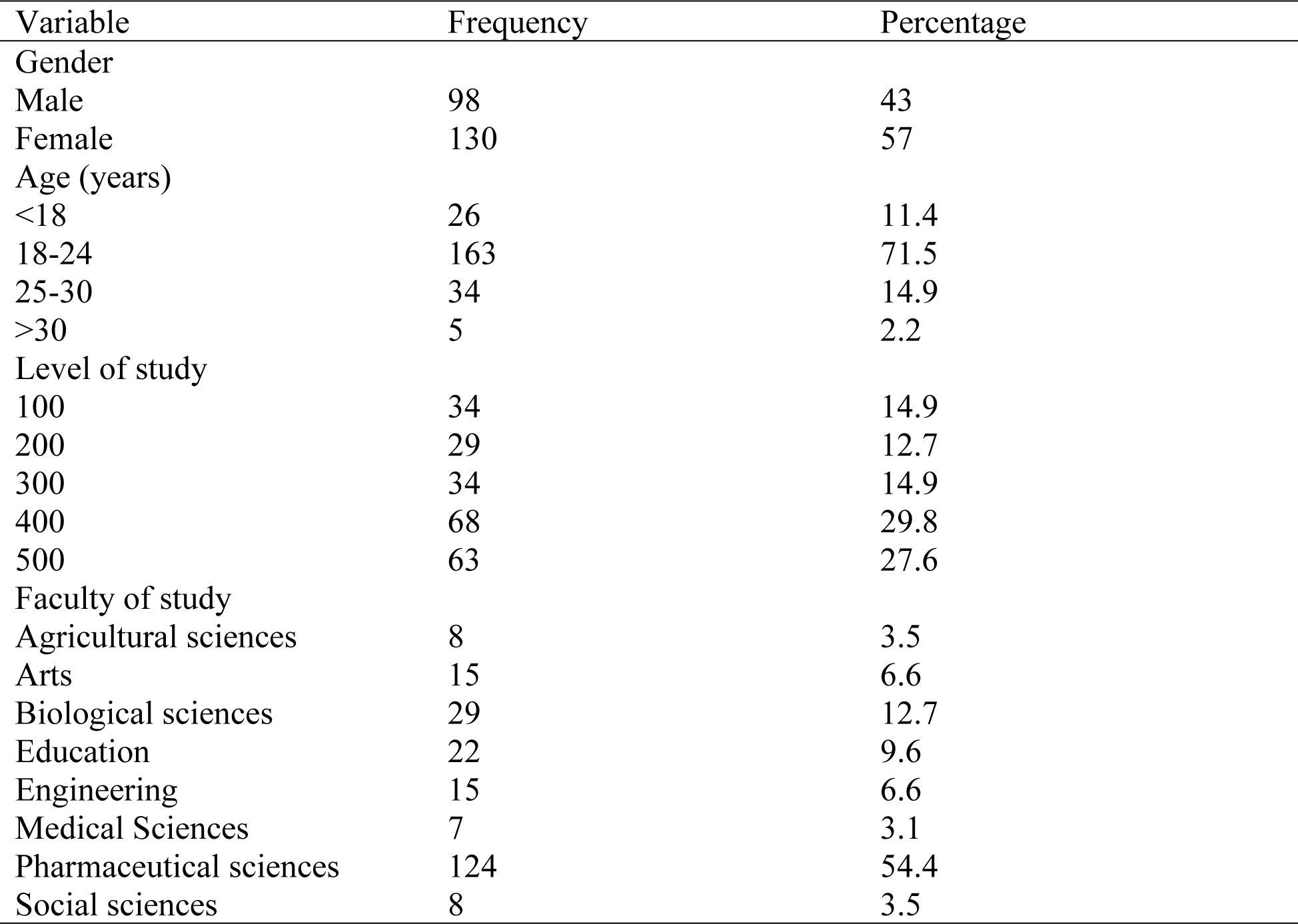
Socio-demographic variables.

**Table 2:** Social Media Utility.

| Social Media Platform | Always<br>N (%) | Very Often<br>N (%) | Sometimes<br>N (%) | Rarely<br>N (%) | Never<br>N (%) | Mean<br>Utility |
| --- | --- | --- | --- | --- | --- | --- |
| WhatsApp | 134 (58.8) | 55 (24.1) | 30 (13.2) | 6 (2.6) | 3 (1.3) | 4.4±0.902 |
| Facebook | 34 (14.9) | 59 (25.9) | 80 (35.1) | 48 (21.1) | 7 (3.1) | 3.3±1.055 |
| Instagram | 26 (11.4) | 33 (14.5) | 70 (30.7) | 46 (20.2) | 53 (23.2) | 2.7±1.285 |
| YouTube | 30 (13.2) | 47 (20.6) | 91 (39.9) | 44 (19.3) | 16 (7.0) | 3.1±1.092 |
| Facebook Messenger | 27 (11.8) | 29 (12.7) | 71 (31.1) | 71 (31.1) | 30 (13.2) | 2.8±1.183 |
| Google Plus | 17 (7.5) | 27 (11.8) | 45 (19.7) | 49 (21.5) | 90 (39.5) | 1.9±1.181 |
| Twitter | 14 (6.1) | 10 (4.4) | 41 (18.0) | 53 (23.2) | 110 (48.2) | 2.3±1.294 |
| SnapChat | 5 (2.2) | 18 (7.9) | 39 (17.1) | 49 (21.5) | 117 (51.3) | 1.9±1.090 |
| Skype | 0 (0) | 2 (0.9) | 15 (6.6) | 47 (20.6) | 164 (71.9) | 1.4±0.646 |
| LinkedIn | 2 (0.9) | 5 (2.2) | 19 (8.3) | 32 (14.0) | 170 (74.6) | 1.4±0.805 |

The mean social media anxiety score was 68.75±12.35. Social media anxiety was higher in females (69.00±12.59) than males (68.42±12.06) although this difference did not reach statistical significance (t = -0.356, p = 0.864). Social media usage was higher in females (35.02±5.04) than males (34.58±6.01) but the difference did not reach statistical significance (t = -0.603, p = 0.314). There was a non-significant negative association between Social media usage and social media anxiety (r = -0.051, p = 0.4450). Social media usage had a negative association with all the domains except Interaction anxiety where association was positive (r = 0.001, p = 0.994). (Table 4). Respondents who had Social media anxiety scores greater than the mean score of 68.75 were categorized as having social media anxiety while those with scores lower than that were regarded as not having social media anxiety. More than half of the students (55.7%, n = 127) had Social media anxiety.

**Table 3:** Social media Anxiety.

| Domains | Maximum | Mean | Standard deviation |
| --- | --- | --- | --- |
| Privacy Concerns | 25 | 19.43 | 3.65 |
| Social Content Anxiety | 35 | 22.5 | 6.02 |
| Self-evaluation | 15 | 10.33 | 2.98 |
| Interaction anxiety | 30 | 16.49 | 5.47 |
| **Total SMA score** |  |  |  |
| Social Media Anxiety | 105 | 68.75 | 12.35 |

**Table 4:** Correlation between Social media usage and social media anxiety.

Correlations
|  |  | SMU | IA | SCA | PCA | SEA | SMA |
| --- | --- | --- | --- | --- | --- | --- | --- |
| SMU | Pearson Correlation | 1 | .001 | -.066 | .017 | -.099 | -.051 |
|  | Sig. (2-tailed) |  | .994 | .323 | .798 | .135 | .445 |
|  | N |  | 228 | 228 | 228 | 228 | 228 |
| IA | Pearson Correlation |  | 1 | .347** | .193** | .214** | .721** |
|  | Sig. (2-tailed) |  |  | .000 | .003 | .001 | .000 |
|  | N |  |  | 228 | 228 | 228 | 228 |
| SCA | Pearson Correlation |  |  | 1 | .236** | .366** | .800** |
|  | Sig. (2-tailed) |  |  |  | .000 | .000 | .000 |
|  | N |  |  |  | 228 | 228 | 228 |
| PCA | Pearson Correlation |  |  |  | 1 | .137* | .529** |
|  | Sig. (2-tailed) |  |  |  |  | .039 | .000 |
|  | N |  |  |  |  | 228 | 228 |
| SEA | Pearson Correlation |  |  |  |  | 1 | .555** |
|  | Sig. (2-tailed) |  |  |  |  |  | .000 |
|  | N |  |  |  |  |  | 228 |
| SMA | Pearson Correlation |  |  |  |  |  | 1 |
|  | Sig. (2-tailed) |  |  |  |  |  |  |
|  | N |  |  |  |  |  |  |

\*\* . Correlation is significant at the 0.01 level (2-tailed).
\* . Correlation is significant at the 0.05 level (2-tailed).
SMU-Social media utility, IA-Interaction anxiety, SCA-Shared content anxiety, PCA- Privacy concern anxiety,
SEA- Self-evaluation anxiety, SMA- Social media anxiety

## Discussion

The ubiquitous use of social media has led to research studies on its psychological implications. This is the first attempt to assess social anxiety resulting from social media use in an African country. The study employed the use of a validated specific questionnaire designed by Alkis et al. to assess social media anxiety among university students. There were more female respondents than male respondents in the present study. This is in contrast to findings from a study to assess the prevalence social anxiety among Nigerian university students where there were more male respondents than female respondents^4^. However, previous research among university students globally has shown response rates to vary by gender with females being more likely to respond than males^14,15^. The students were mostly within the age 18 and 24 years and were in their fourth year of study. This is in line with the 6-3-3-4 system of education obtainable in Nigeria where the minimum age of entry into public universities is pegged at 16 years. The 6-3-3-4 system started in 1987 after the National Policy on Education was introduced^16^. This policy was initiated to unify the structure of education all over the country.

Globally, the use of social media has steadily been on the increase and the most frequent users are reported to be young adults between 18-29 years ^17,18^. The most frequently used social media platform in this study were WhatsApp and Facebook. This finding concurs with current literature on the use of social media among students in Nigerian tertiary institutions^19,20^. Skype and LinkedIn were the least frequently used social media networks probably because the respondents were students who were yet to build a career and may be less concerned about building a professional profile. Social media usage was higher among females. This is in concordance with available data on gender differences in social media use^21^. Social media anxiety were higher among females than males. This concurs with findings from the Nigerian study on social phobia among university students where prevalence was found to be higher among females^4^.

In the current study, there was a non-significant negative association between Social media usage and social media anxiety. This finding may be explained by the social enhancement hypothesis which proposes that socially skilled individuals use online social platforms to increase their chances of interacting with others^22^. The study findings agrees with findings from studies carried out Rizvi among Pakistani students where no significant relationship was found between social anxiety and Facebook usage^23^. Another study by Kross and his colleagues found no relationship between Facebook use frequency and ratings of worry in a small sample of young adults^24^ On the other hand studies across the globe have found significant positive associations between social media usage and social anxiety^25,26^. This positive correlation may be explained by the social compensation hypothesis which proposes that individuals use online social networking sites to compensate for deficits in social skills or discomfort in face-to-face situations^22^.

Social media usage had a non-significant positive association with Interaction anxiety in this study. A study by Henry among university students found that psychological stressors became manifest when they relied on social media to cope with personal problems indicating a lack of ability to build healthy interpersonal relationships^27^. A meta-analysis by Shechner, and Aderka found a positive correlation between social anxiety and feeling comfortable online^28^. Privacy concern, Shared content anxiety and self-evaluation anxiety had non-significant negative associations with social media usage. Research has shown that individuals with social anxiety perceive online interactions to be less threatening than real-life conversations and thus tend to self-disclose more in online settings^29,30^.

In conclusion, there was a negative non-significant correlation between social media usage and social anxiety. Whatsapp was the most frequently used social media platform among the university students surveyed. This study has contributed to the sparse literature available on social anxiety in social media. Investigations regarding this issue are scarce from low and middle income countries and this is the first from an African country. Additional strengths include the use of a standard well-validated measure specific for social media anxiety. However, the cross-sectional design may limit the conclusions about causality. Another limitation is the self-reporting nature of the questionnaire used to assess social media use and anxiety. Longitudinal studies are necessary to investigate whether social media use is a causal risk factor for anxiety symptoms and anxiety-related disorders and to assess the impact of anxiety on social media use.

## Acknowledgement

The authors wish to acknowledge the role of members of the International Society for Pharmacoeconomics and Outcomes Research group (ISPOR) University of Nigeria Nsukka in the data collection phase of this study.

